# Classification of Brain Tumor IDH Status using MRI and Deep Learning

**DOI:** 10.1101/757344

**Authors:** Sahil Nalawade, Gowtham Murugesan, Maryam Vejdani-Jahromi, Ryan A. Fisicaro, Chandan Ganesh Bangalore Yogananda, Ben Wagner, Bruce Mickey, Elizabeth Maher, Marco C. Pinho, Baowei Fei, Ananth J. Madhuranthakam, Joseph A. Maldjian

**Affiliations:** Department of Radiology, UT Southwestern Medical Center, Dallas, TX, USA, 75235; Department of Neurological Surgery, UT Southwestern Medical Center, Dallas, TX, USA, 75390; Department of Neurology & Neurotherapeutics, UT Southwestern Medical Center, Dallas, TX, USA, 75390; Department of Bioengineering, UT Dallas, Richardson, TX, USA, 75080

**Author notes:** Equal Contribution (Sahil Nalawade, Gowtham Murugesan).

**Keywords:** Isocitrate dehydrogenase (IDH), MRI, Convolutional networks, Deep learning, Tumor classification, Radiomics

## Abstract

Isocitrate dehydrogenase (IDH) mutation status is an important marker in glioma diagnosis and therapy. We propose a novel automated pipeline for predicting IDH status noninvasively using deep learning and T2-weighted (T2w) MR images with minimal preprocessing (N4 bias correction and normalization to zero mean and unit variance). T2w MRI and genomic data were obtained from The Cancer Imaging Archive dataset (TCIA) for 260 subjects (120 High grade and 140 Low grade gliomas). A fully automated 2D densely connected model was trained to classify IDH mutation status on 208 subjects and tested on another held-out set of 52 subjects, using 5-fold cross validation. Data leakage was avoided by ensuring subject separation during the slice-wise randomization. Mean classification accuracy of 90.5% was achieved for each axial slice in predicting the three classes of no tumor, IDH mutated and IDH wild-type. Test accuracy of 83.8% was achieved in predicting IDH mutation status for individual subjects on the test dataset of 52 subjects. We demonstrate a deep learning method to predict IDH mutation status using T2w MRI alone. Radiologic imaging studies using deep learning methods must address data leakage (subject duplication) in the randomization process to avoid upward bias in the reported classification accuracy.

## 1 Introduction

In 2008 it was reported that some glioblastomas harbor a mutation in a gene coding for the citric acid cycle enzyme isocitrate dehydrogenase (IDH) (1). Subsequent studies revealed that the majority of low grade gliomas possess a mutant form of IDH, and that the mutant enzyme catalyzes the production of the oncometabolite 2-hydroxyglutarate (2-HG) (2). Although this product of the mutant form of IDH is believed to play a role in the initiation of the neoplastic process, it has been observed that gliomas that contain the mutant enzyme have a better prognosis than tumors of the same grade that contain only the wild type IDH. This observation implies that IDH mutated and IDH wild type gliomas are biologically different tumors, and led the World Health Organization (WHO) to designate them as such in the latest revision of their classification of gliomas (3). Although a presumptive diagnosis of an IDH mutated glioma may be made on the basis of MR spectroscopy for 2-HG (4–7), at the present time, the only way to definitively identify an IDH mutated glioma is to perform immunohistochemistry or gene sequencing on a tissue specimen, acquired through biopsy or surgery. Because the differences between IDH mutated and IDH wild type gliomas may have implications for their treatment, especially if inhibitors of the mutant IDH enzyme currently in development prove to halt their growth, there is interest in attempting to distinguish between these two tumor types prior to surgery. As noted above, one avenue of research involves using MR spectroscopy to measure levels of 2-HG in the tumor (5, 8–10). More recent studies have attempted to utilize machine learning techniques to analyze diagnostic MR images and predict IDH mutation status in gliomas using anatomic differences between the two tumor types.

Delfanti et al. demonstrated that genomic information with fluid attenuated inversion recovery (FLAIR) MRI could be used for the classification of patient images into IDH wild type, and IDH mutation with and without 1p/19q co-deletion (11). The main determinants for classification were tumor border and location, with IDH mutant tumors having well-defined or slightly ill-defined borders and predominantly a frontal localization; and IDH wild type tumors demonstrating undefined borders and location in non-frontal areas. Chang et al. developed a deep learning residual network model for predicting IDH mutation with preprocessing steps including resampling, co-registration of multiple sequences, bias correction, normalization and tumor segmentation (12). Using a combination of imaging and age, the model demonstrated testing accuracy of 89.1% and an area under the curve (AUC) value of 0.95 for IDH mutation for all image sequences combined. Zhang et al. used 103 low grade glioma (LGG) subjects for training a support vector machine (SVM) for classifying IDH mutation status, achieving an AUC of 0.83 on testing data (13). In another approach, Chang et al (14) similarly demonstrated that IDH mutation status can be determined using T2-weighted (T2w), T2w- Fluid attenuated inversion recovery (FLAIR) and T1-weighted pre- and post-contrast images. Preprocessing steps in their work included co-registration of all sequences, intensity normalization using zero mean and unit variance, application of a 3D convolutional neural network (CNN) based whole tumor segmentation tool for segmenting the lesion margins, cropping the output tumor mask on all input imaging sequences, and resizing individual image slices to 32 × 32 with 4 input sequence channels. The mean accuracy result from the model was 94% with a 5-fold cross validation accuracy ranging from 90% to 96% (14). Common to all of these previous methods is the involvement of preprocessing steps, typically including some form of brain tumor pre-segmentation or region of interest extraction, and utilizing multiparametric or 3D near-isotropic MRI data that is often not part of the standard clinical imaging protocol (12, 14).

In this work, we propose a fully automated deep learning based pipeline using a densely connected network model, that involves minimal preprocessing and requires only standard T2w images. A similar approach has been previously used for the identification of the O^6^ – methylguanine-DNA methyltransferase (MGMT) methylation status and prediction of 1p/19q chromosomal arm deletion (15). Clinical T2-weighted images are acquired in a short time frame (typically around 2 minutes), and are robust to motion with current acquisition methods. Almost universally, high quality T2-weighted images are acquired during clinical brain tumor work-ups. The preprocessing steps preserve the original image information without the need for any resampling, skull stripping, region-of-interest, or tumor pre-segmentation procedures. The advantage of a dense network model is that it passes the weights from all the previous blocks to the subsequent blocks, preserving the information from the initial layer and aiding in the classification.

The ability to quickly and accurately classify IDH status non-invasively can help with better planning, counseling, and treatment of brain tumor patients, especially in cases where biopsy is not feasible due to unfavorable tumor locations. A methodologic contribution that we make specifically to the radiologic deep learning literature is on the approach to data randomization for 2D models. Furthermore, the deep learning approach is fully automated and can be easily implemented in the clinical workflow using only T2-weighted MR images.

## 2 Materials and Methods

### 2.1 Subjects

260 subjects from The Cancer Imaging Archive (TCIA) (16) dataset were selected, including 120 high grade gliomas (HGG) (17) and 140 low grade gliomas (LGG) (18), and based on their pre-operative status from a pool of 461 subjects. The genomic information was provided through the National Cancer Institute - Genomic Data Commons (GDC) Data Portal (19). The genomic data was available in the following 3 classes: IDH mutated, IDH wild type, and Not Available (N/A). The Genomic data of the N/A type was excluded from the pool of 461 subjects. MRI data was filtered for any visible artifacts in the images. The final dataset consisted of 260 subjects based on the available genomic information, MRI data, pre-operative status and lack of image artifacts on the T2w images.

A standard 80:20 data split was employed with 80% training and 20% testing (held-out). The 80% training was further split into a standard 80:20 split of 80% training and 20% validation. The final dataset of 260 subjects was thus randomly divided into a training set (208 subjects, including approximately 96 HGG and 112 LGG) and a test set (52 subjects, including approximately 24 HGG and 28 LGG). This process was repeated separately for each fold during the 5-fold cross validation.

For each fold of the cross-validation, 208 subjects with, on average, 9,728 axial slices of T2w images were selected for training and validation (7177 slices – No tumor, 1110 slices – IDH mutated, 1441 slices– IDH wild type). The start and end slices of the tumor (edge slices) were manually labeled for each T2 dataset. These edge slices were excluded from training to provide more robust ground truth data. All slices were included for the testing set. Each T2w slice was manually assigned only one label (No Tumor, IDH mutated, or IDH wild type). In order to address any class imbalance due to the higher number of no tumor slices, class weights were assigned based on the labels in the training dataset. Although this was a slice-wise training model, slices of subjects in the testing set were not mixed into the training set. This is a critical step related to the data leakage problem in 2D networks, especially for radiologic deep learning studies (20, 21). This was necessary to avoid bias during testing and an over inflation of the measured accuracies. Fifty two subjects with 2522 axial slices (1839 slices– No Tumor, 299 slices– IDH mutated, 384 slices– IDH wild type) were not included in the training or validation and were used for testing, for each fold. Classification was done on a slice-wise basis (2D) followed by majority voting across all slices to provide a patient-level classification. Note that we use the term slice-wise to refer to classification of each 2D axial image for IDH status. Similarly, the term subject-wise is used for classification of IDH status for each subject. We used a straightforward majority voting scheme to determine subject-wise classification based on the majority IDH classification of the individual 2D slices. Subjects classified with an equal number of IDH mutated and IDH wild-type tumor slices were assigned to the IDH wild type group.

### 2.2 Image Processing

Minimal standard preprocessing of the T2w images from the TCIA data set was performed prior to training (Figure 1). The images were converted from DICOM to nifti format using dcm2nii, bias corrected to remove RF inhomogeneity using the N4 bias correction algorithm, zero-mean intensity normalized to between −1 and 1, and resampled to 128 × 128 image dimensions to improve the computational efficiency during training. The Inception V4 model however, required input image size of 299 × 299 as a design constraint of this model when originally constructed (22, 23). The total preprocessing time for each subject was less than 1 minute.

**Fig 1.**
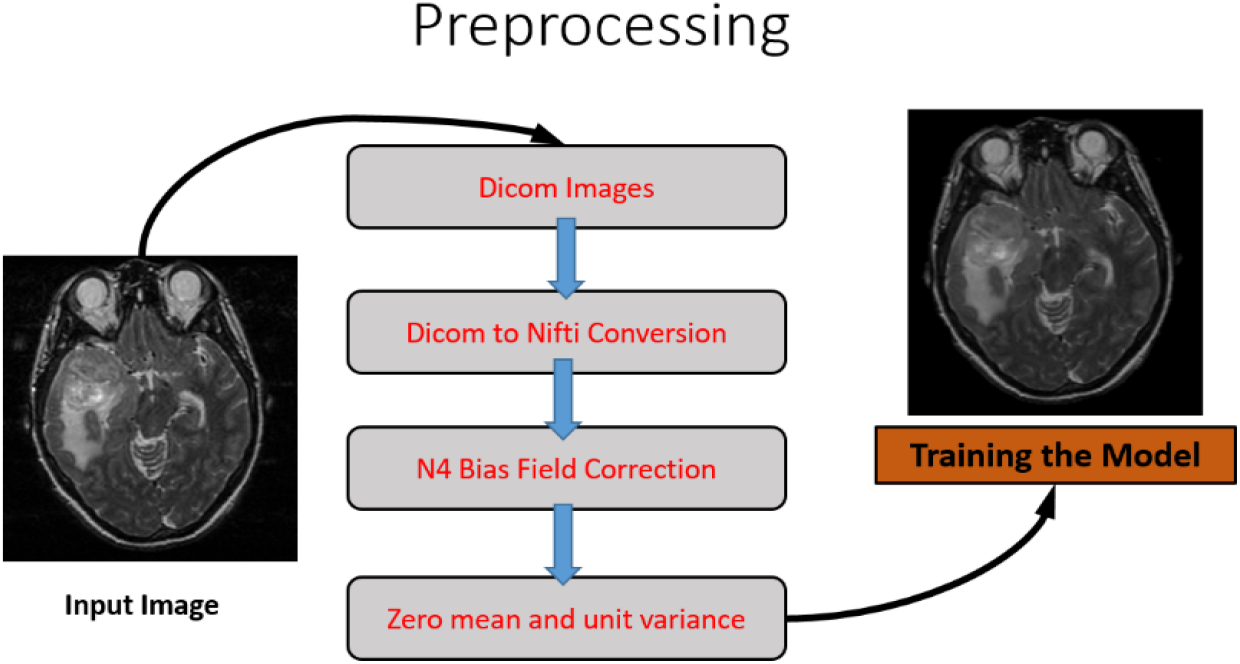
Flowchart of preprocessing steps prior to training the deep learning model

### 2.3 Model Training

The following models were used for classification of the T2w images into IDH mutated and IDH wild type classes: residual network (ResNet-50), Densely connected network (DenseNet-161), and Inception-v4. Our choice of network architectures was based on the best performers from the ImageNet challenge for 2015 (ResNET), and 2017 (DenseNet and Inception V4). The DenseNet model, designed by Huang *et al* (24) received the best paper award at CVPR 2017. The models were trained with the Pycharm and Python IDEs using the Keras python package with TensorFlow backend engines. Fine tuning of the 3 classes was performed on all models. The three-class labels for each slice were: no tumor, IDH mutated, and IDH wild type. The models were originally trained on ImageNet data with 3 channels (RGB). For our implementation the 3-channel input was provided as a central slice with the 2 immediate surrounding slices. If the central slice was the first or last slice, the surrounding slices were assigned as no value.

### 2.4 ResNet-50 Model

The residual network was implemented as proposed by He et al. (25). Each residual connection adds the input of the block to the output, helping to preserve the information from the previous block. A deep residual network framework was added to the model while maintaining parameter numbers to address issues with convergence in the originally proposed model. The residual net used the kernel initializer as ‘He normal’ for weight initialization. On top of the residual network model, a flattened output was added and sent to the dense layer with the rectified linear unit (‘relu’) activation and a dropout of 0.5. The final layer of the model was the classification layer with a softmax activation and the number of classes as the output. The residual network model used for training was ResNet-50 (Figure 2).

**Fig. 2.**
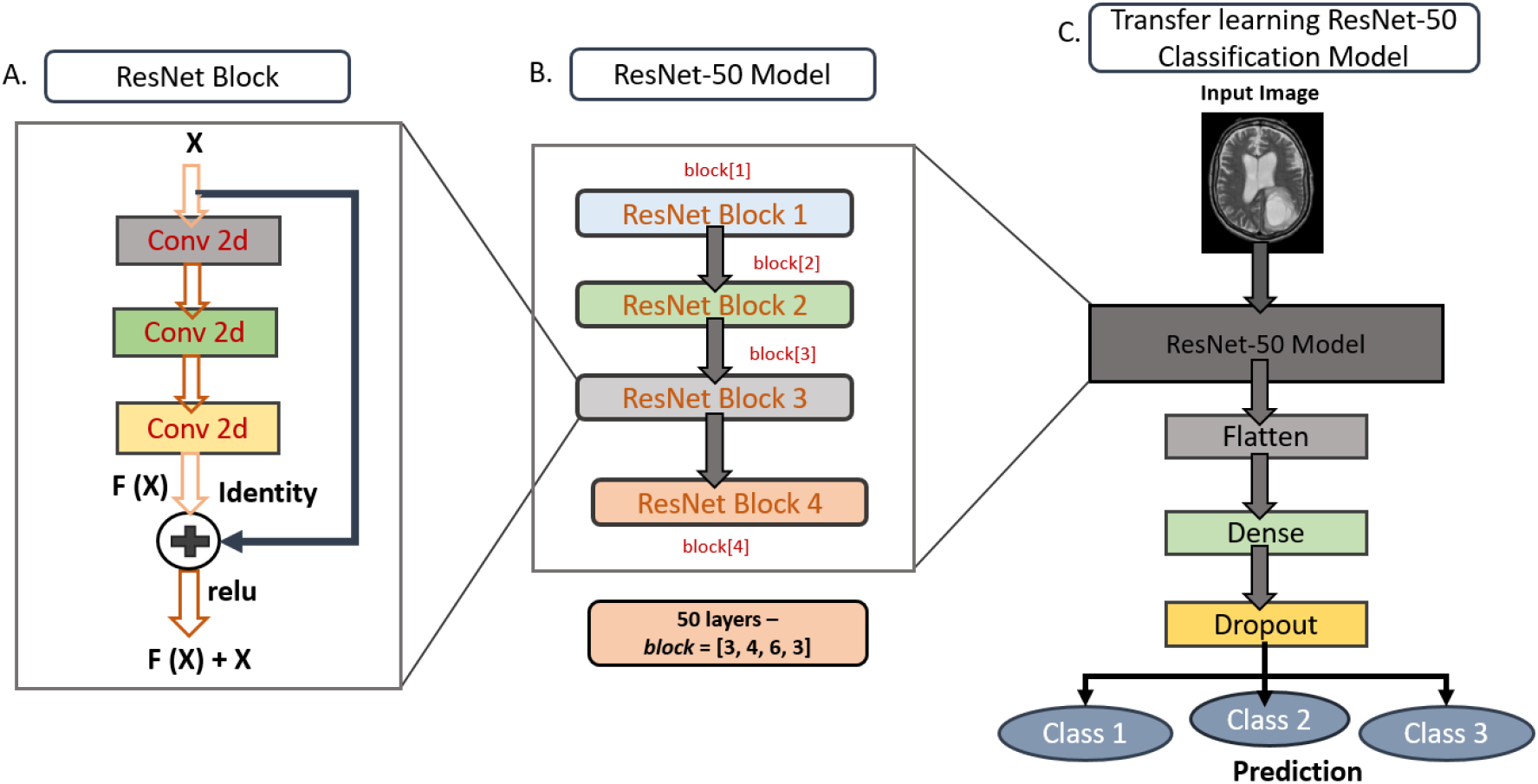
Architecture of ResNet-50 (50 layers) Model used for IDH classification

### 2.5 Inception-v4 Model

The Inception model architecture was designed by the Google Research team (22, 23). The Inception-v4 model is a deep architecture with 41 million parameters and the model is designed with inception blocks and reduction blocks. The inception blocks are used in a sequential manner with reduction blocks except for the last inception block, which has an average pooling layer and a dropout layer before the classification layer.

### 2.6 DenseNet-161 Model

The DenseNet model was based on the design by Huang et al. (24). This model was inspired by the residual network model, which allows the residual connections to pass information from the previous layer to the subsequent layer. Dense networks have advantages over other networks by alleviating the vanishing gradient problem with feature propagation through the dense connection to the subsequent layers.

The features passed to the subsequent layers in the DenseNet model are not added by summation, but are combined using concatenation. Each block has connections from the previous block such that L=number of blocks and the number of connections for each block is L×(L+1)/2, creating a dense connectivity pattern or DenseNet. The DenseNet-161 model architecture is shown in Figure 3, which illustrates a 5 block approach where the 1^st^ block is the Input layer and each of the subsequent 4 blocks are characterized by 2D convolution layers with filter size of (1 × 1) and (3 × 3) respectively. The pre-trained model was used to transfer learning and used for classification based on the trained information. A 161 layer DenseNet model was used for model training.

**Fig 3.**
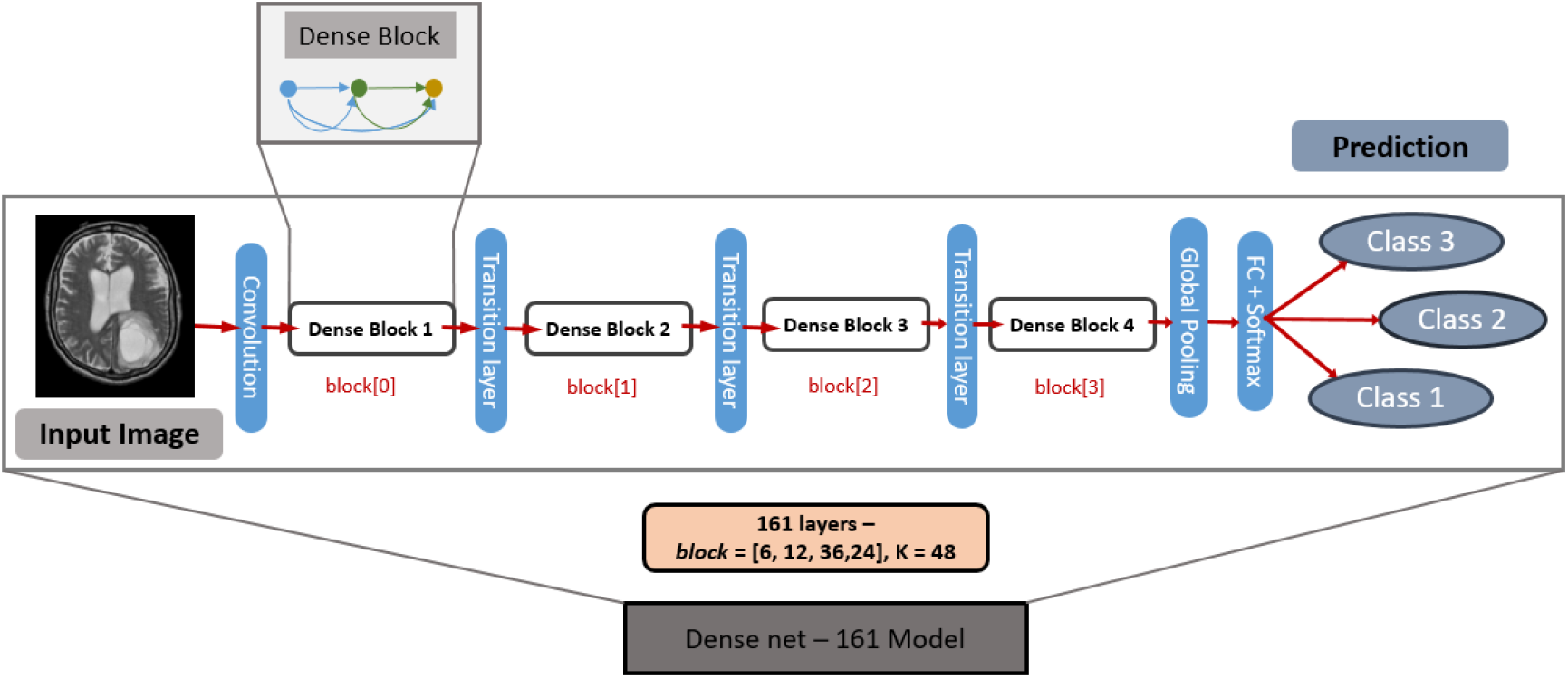
Architecture of the DenseNet-161 (161 layers) Model used for IDH classification.

### 2.7 Training, Testing and Statistical Analysis

Model training was performed on a Nvidia Tesla P100, P40, K40/K80 GPU with 384 GB RAM and the model accuracy was assessed for 200 Epochs. The optimizer used for training was the Stochastic Gradient Descent (26) as described in Zhang et al.(27) and the learning rate was set to 10^−5^, with a decay of 10^−7^ and momentum of 0.8. Data augmentation was performed on the training dataset, which included vertical and horizontal flip, random rotation, translation, shear, zoom shifts and elastic transformation to minimize overfitting the data. The results were analyzed by assessing accuracy, precision, sensitivity, specificity, and F-1 score values. Figure 4 shows the confusion matrix and the equations for calculating the testing parameters. Slice-wise model testing was performed based on the output from the 2D model. Subject-wise classification was performed based on majority voting across IDH mutated and IDH wild type tumor slices. This classification accuracy was computed on the independent test dataset that was separate from the testing and validation data sets.

**Fig 4.**
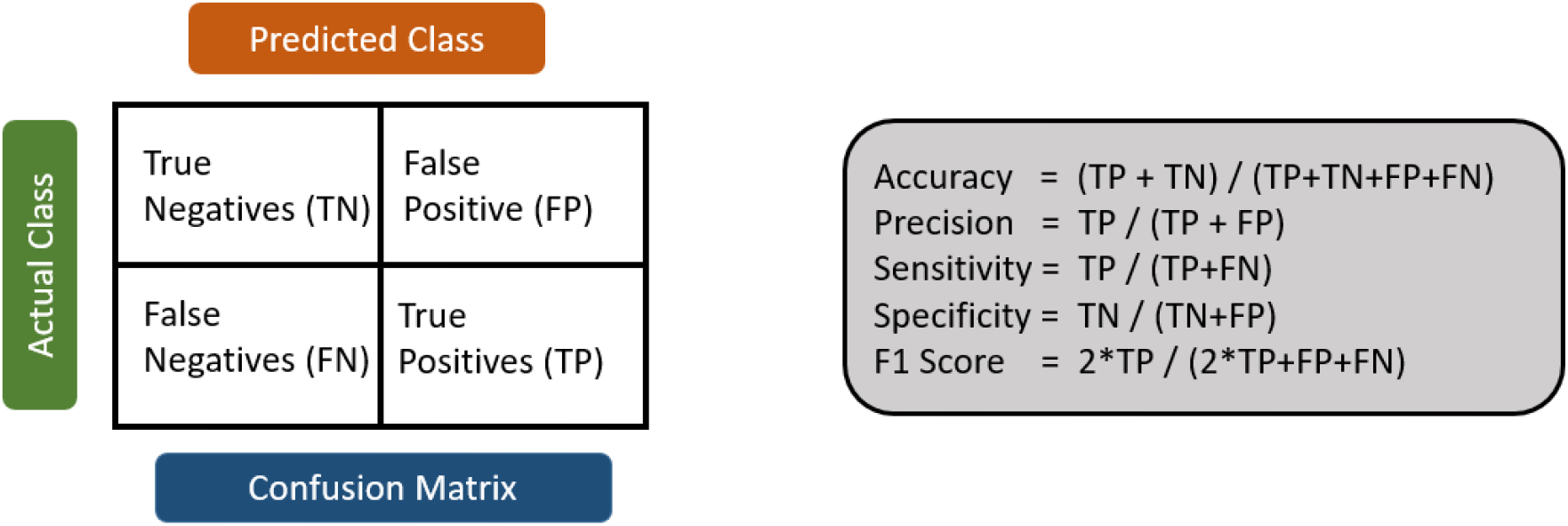
Confusion Matrix and equations for calculating accuracy, precision, sensitivity, specificity, and F1 score

### 2.8 Model training times

The DensetNet-161 model took approximately 110 hours for training, while the ResNet-50 model and the Inception V4 model took approximately 56 hours and 32 hours, respectively. Testing time for individual subject classification was less than 30 seconds for all models.

## 3 Results

### 3.1 Training, validation, and testing accuracy

Table 1 shows the accuracy comparison between the ResNet-50, DenseNet-161 and Inception-v4 models. The DenseNet-161 model was superior in training, validation, and testing accuracy compared to ResNet-50 and Inception-v4. Averaged across the five folds, the slice-wise accuracy of the DenseNet-161 model was 90.5 ±1.0% (standard deviation) with an AUC of 0.95 on the held out test dataset of 52 subjects.

**Table 1.** Slice-wise accuracy comparisons between the ResNet-50 Model, Inception-v4, and DenseNet-161 model averaged for 5 fold cross validation

| ***Results averaged for 5 fold cross validation*** |  |  |  |
| --- | --- | --- | --- |
| **Model** | **Training accuracy (%)** | **Validation accuracy (%)** | **Testing accuracy (%)** |
| Inception-v4 | 64.8 | 72.2 | 1916/2522 (76.1) |
| ResNet-50 | 97.9 | 96.5 | 2265/2522 (89.7) |
| **DenseNet-161** | **97.9** | **96.4** | **2282/2522 (90.5)** |

### 3.2 Accuracy, Precision, Recall/Sensitivity, Specificity, F1 score and AUC Comparison

Average metrics were computed across folds and classes. The classification accuracy, precision, recall/sensitivity, specificity, F1 score and AUC for slice-wise IDH classification with the DenseNet-161 model were 90.5 ±1.0%, 79.9 ±3.4%, 83.1 ±3.2%, 94.8 ±0.5%, 81.3 ±3.2% and 0.95, respectively. For subject-wise IDH classification, accuracy, precision/positive predictive value, recall/sensitivity, specificity, F1 score and AUC were 84.1 ±2.9%, 83.5 ±3.5%, 83.5 ±3.5%, 83.5 ±3.1%, and 0.84 (Table 2). Slice-wise and subject-wise comparisons of accuracy, precision, recall/sensitivity, specificity, F1 score and AUC for each of the 5 fold cross validations for the DenseNet-161 model are shown in Table 3.

**Table 2.**
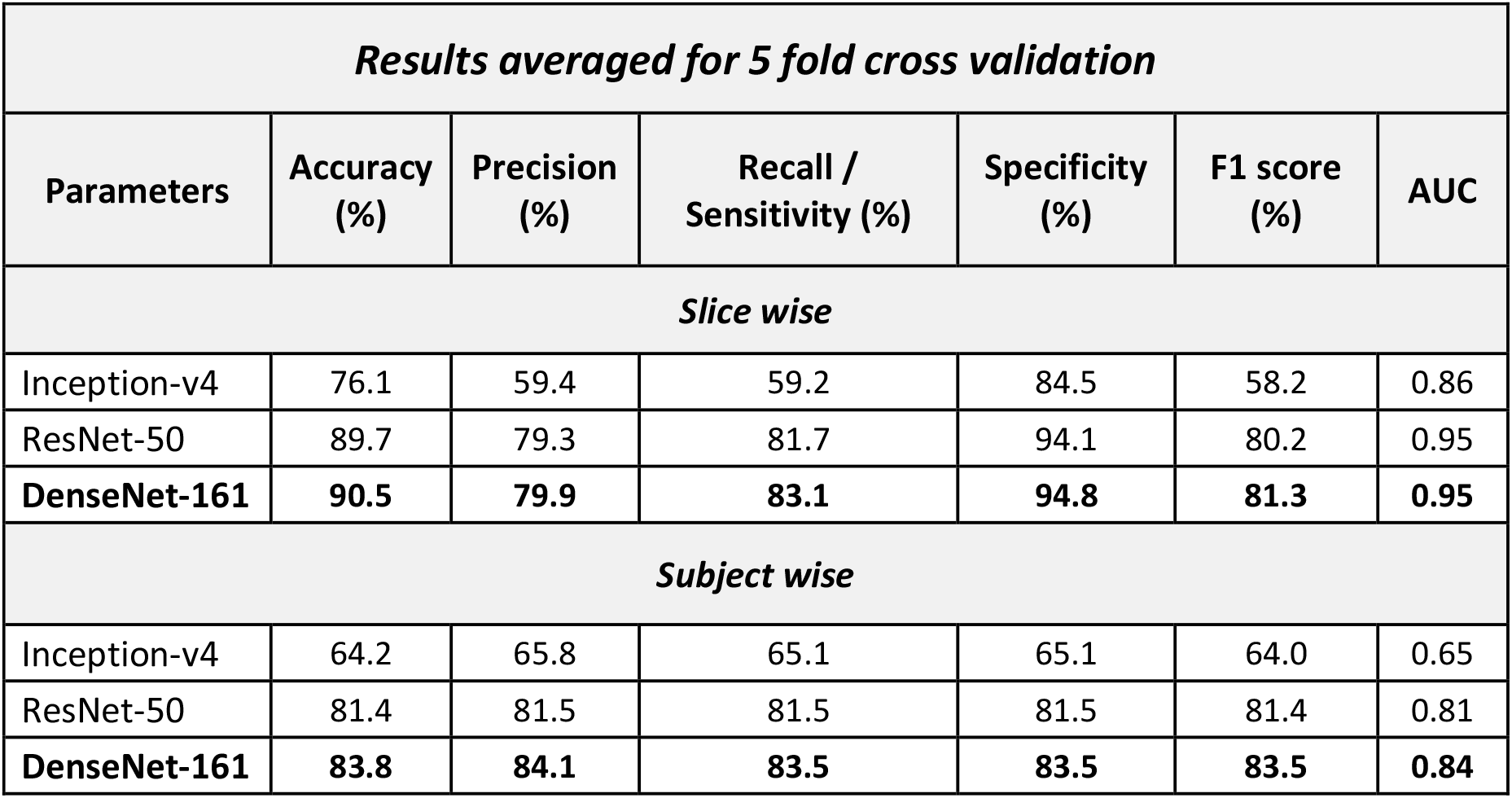
Slice-wise and subject-wise comparison of accuracy, precision, recall, F1 score and AUC parameters averaged for 5 fold cross validation for ResNet-50, Inception-v4 and DenseNet-161

**Table 3.** Slice-wise and subject-wise comparison of accuracy, precision, recall, F1 score and AUC parameters for each of the 5 fold cross validations for the DenseNet-161 model.

| ***DenseNet-161 model*** |  |  |  |  |  |  |
| --- | --- | --- | --- | --- | --- | --- |
| **Fold** | **Accuracy (%)** | **Precision (%)** | **Recall / Sensitivity (%)** | **Specificity (%)** | **F1 score (%)** | **AUC** |
| ***Slice wise*** |  |  |  |  |  |  |
| 1 | 91.7 | 83.3 | 86.0 | 94.9 | 84.4 | 0.95 |
| 2 | 91.0 | 82.7 | 84.4 | 95.1 | 83.5 | 0.95 |
| 3 | 90.1 | 79.3 | 81.5 | 94.5 | 80.2 | 0.95 |
| 4 | 88.7 | 73.7 | 77.6 | 93.9 | 75.5 | 0.91 |
| 5 | 90.9 | 80.3 | 86.0 | 95.4 | 82.8 | 0.95 |
| ***Subject wise*** |  |  |  |  |  |  |
| 1 | 84.6 | 84.8 | 84.0 | 84.0 | 84.2 | 0.84 |
| 2 | 86.5 | 87.2 | 87.5 | 87.5 | 86.5 | 0.87 |
| 3 | 78.8 | 79.1 | 77.9 | 77.9 | 78.2 | 0.78 |
| 4 | 82.7 | 83.1 | 81.8 | 81.8 | 82.2 | 0.82 |
| 5 | 86.5 | 86.3 | 86.6 | 86.6 | 86.4 | 0.87 |

### 3.3 Slice-wise comparison

The precision for the DenseNet-161 model across 5 fold cross validation was 97.7 ±0.5% for the “no tumor” classification, 71.7 ±6.8% for IDH mutation, and 70.3 ±5.5% for IDH wild type.

### 3.4 Subject-wise comparison

For the DenseNet-161 model, the sensitivity and specificity for subject-wise IDH mutation classification was 80.9 ±9.4% and 86.2 ±3.8%, respectively. The positive and negative predictive values were 82.5 ±2.8% and 85.7 ±6.3%, respectively.

## 4 Discussion

The results from Tables 1 and 2 show that the ResNet-50 model performed better than the Inception-v4 model. The ResNet-50 architecture has residual connections which preserve information from the previous layer in the residual block. The DenseNet-161 model performed the best of all the three models tested. Unlike the ResNet-50 model, the DenseNet-161 model architecture carries the information from all previous layers and adds the information to the next layer. This helped in learning the information from different layers and transferring to the next layers. The slice-wise classification AUC results were 0.95 for DenseNet-161, 0.95 for ResNet-50, and 0.86 for Inception-v4.

Chang et al. (14) demonstrated a high classification accuracy for IDH mutation status using T2w, FLAIR, T1w pre- and post-contrast images. Preprocessing steps included coregistration across multiple sequences, intensity normalization to zero mean and unit variance, segmentation of the brain tumor and cropping the images and resizing slices to 32 × 32. A 94% mean accuracy on 5-fold cross validation was reported. The approach to classification was slice-wise, similar to our model. In designing the slice-wise classification model, it is important to ensure that none of the slices of subjects from the testing set are inadvertently included in the training set. This can easily be overlooked in 2D slice-wise models during the slice randomization process that generate the training slices, validation slices, and testing slices. This can introduce bias in the testing phase artificially boosting accuracies by including slices from subjects in the training set that share considerable information with different slices but from the same subjects in the testing set. It is not clear in the previously reported 2D models whether this caveat was adhered to.

An important methodologic contribution that we make specifically to the radiologic deep learning literature is on the approach to data randomization for 2D models. It is critical that imaging researchers are aware of the data leakage and subject duplication issue. This is perhaps unique to radiology where multiple slices of pathology are acquired in MRI or CT, with considerable overlap in feature content from slice to slice. Widely used deep learning tools provide the ability to perform data randomization using a simple flag in the called routine (*e.g.*, in Keras, or Scikit-learn(28)). Use of this flag in 2D imaging based CNNs can lead to bias in the results by inadvertently including slices from the same subject in both training and testing cohorts. This is a significant concern, as it can lead to data leakage in which examples of the same subject (albeit different slices of the same tumor) can appear in the training set and the test set. The problem of data leakage in medical images was discussed by Wegmayr et. al. (29) and Feng et. al.(20) and has been referred to as subject duplication in training and testing sets. In our initial studies, we did not account for the data leakage problem and achieved accuracies of 95% with the T2 images alone, slightly higher than that of Chang et al (14). When appropriately accounting for the data leakage issue, our accuracies were reduced to the 83.8% reported here. One of the major contributions of our work is in making the radiology community aware of the data leakage problem, as it is very easy to overlook when 2D networks are considered that use image slices as input.

The majority of HGG tumors are IDH wild type (up to 90%). An algorithm that merely distinguishes between HGG and LGG for determination of IDH status is likely of limited value as this can be done subjectively with fairly high accuracy on the basis of contrast enhancement. For example, previous studies that used multiparametric MR data for determination of IDH status in HGG and LGG may have demonstrated high accuracy predominantly on the basis of contrast enhancement features. The more valuable distinction from a clinical standpoint would be between IDH mutated and IDH wild type low grade gliomas in which contrast enhancement is usually absent. Our training and testing samples were weighted towards LGG, and there were a significant number of IDH wild type LGG in both the training and validation sample (~ 30%). Our testing accuracy for the LGG group was 78.6%. Additionally, our use of T2w-only images eliminates the potential for the algorithm being a contrast-enhancement discriminator.

Our method provides high accuracy with minimal preprocessing steps as compared to previous work. The preprocessing steps in our work only involve N4 bias field correction and intensity normalization. Our method also involves no tumor segmentation or ROI extraction as described in Chang et al.(12), which helps in reducing the time, effort and potential sources of error. Our method also does not require pre-engineered features to be extracted from the images or histopathological data as described in Delfanti et al. (11). This general approach can be easily incorporated into an automated clinical workflow for IDH classification. The minimal preprocessing, and the use of standard T2w images alone makes it promising as a robust clinical tool for noninvasively determining IDH mutation status.

## 5 Limitations

This is a retrospective study applying several neural network architectures to the TCIA HGG-LGG database to generate a model predicting IDH genotype based only on T2-weighted MR imaging. The data set, especially at the subject level, is small in terms of deep learning applications and may not generalize well. Fluctuation of performance is also a concern with small data sets. However, the TCIA dataset is the largest curated brain tumor dataset publicly available, and it uses data from multiple sites using different imaging protocols. This database consisted of data from 10 different institutions out of which 8 institutions contributed GBM/HGG datasets and 5 institutions contributed LGG datasets to the TCIA cohort. This provided a very heterogeneous dataset, and we believe this is perhaps even better than using data from a single source for deep learning applications. While our current study focused on the classification of T2w images into no tumor, IDH mutated, an IDH wild type, future studies can extend this approach to classify IDH1 and IDH2 subtypes. Accuracies may be further improved with the inclusion of multiparametric imaging data in the training model. Our approach, however, is much more straightforward using T2-weighted images alone without the requirement of additional imaging sequences. Clinically, T2-weighted images are typically acquired within 2 minutes, and are robust to patient motion. The multi-sequence input required by previous approaches can be compromised due to patient motion from lengthier examination times, and the need for gadolinium contrast, especially as the post-contrast images are typically acquired at the end of an already lengthy examination time. For a potential clinical solution, the use of T2-weighted images is a significant strength, as these images are almost uniformly acquired without artifacts from patient motion.

## 6 Conclusion

We demonstrate a deep learning method to predict IDH mutation status using T2-weighted MR images alone. The proposed model requires minimal preprocessing to obtain high accuracies, without the need for tumor segmentation or extraction of regions of interest, making it promising for robust clinical implementation.

## Acknowledgments

Support for this research was provided by NCI U01CA207091 (AJM, JAM).

