## Supplemental Table 1 for "Classification of Brain Tumor IDH Status using MRI and Deep Learning"

| Subject No. | Subject ID | IDH Status |
|---|---|---|
| 1 | 'TCGA-06-0128' | IDH mutated |
| 2 | 'TCGA-06-0130' | IDH wild type |
| 3 | 'TCGA-06-0137' | IDH wild type |
| 4 | 'TCGA-FG-6689' | IDH mutated |
| 5 | 'TCGA-FG-6690' | IDH mutated |
| 6 | 'TCGA-FG-6691' | IDH mutated |
| 7 | 'TCGA-06-0184' | IDH wild type |
| 8 | 'TCGA-HT-A5R5' | IDH mutated |
| 9 | 'TCGA-06-0188' | IDH wild type |
| 10 | 'TCGA-02-0046' | IDH wild type |
| 11 | 'TCGA-02-0047' | IDH wild type |
| 12 | 'TCGA-06-0160' | IDH wild type |
| 13 | 'TCGA-02-0048' | IDH wild type |
| 14 | 'TCGA-06-0185' | IDH wild type |
| 15 | 'TCGA-06-6389' | IDH mutated |
| 16 | 'TCGA-08-0244' | IDH wild type |
| 17 | 'TCGA-08-0348' | IDH wild type |
| 18 | 'TCGA-08-0350' | IDH wild type |
| 19 | 'TCGA-08-0353' | IDH wild type |
| 20 | 'TCGA-08-0355' | IDH wild type |
| 21 | 'TCGA-08-0356' | IDH wild type |
| 22 | 'TCGA-CS-6666' | IDH mutated |
| 23 | 'TCGA-CS-6667' | IDH mutated |
| 24 | 'TCGA-HT-7467' | IDH mutated |
| 25 | 'TCGA-HT-7616' | IDH mutated |
| 26 | 'TCGA-HT-7602' | IDH mutated |
| 27 | 'TCGA-02-0033' | IDH wild type |
| 28 | 'TCGA-02-0034' | IDH wild type |
| 29 | 'TCGA-02-0037' | IDH wild type |
| 30 | 'TCGA-HT-8563' | IDH mutated |
| 31 | 'TCGA-HT-7884' | IDH mutated |
| 32 | 'TCGA-HT-7874' | IDH mutated |
| 33 | 'TCGA-08-0359' | IDH wild type |
| 34 | 'TCGA-08-0380' | IDH wild type |
| 35 | 'TCGA-08-0389' | IDH wild type |
| 36 | 'TCGA-08-0390' | IDH wild type |
| 37 | 'TCGA-12-0616' | IDH wild type |
| 38 | 'TCGA-12-1093' | IDH wild type |
| 39 | 'TCGA-14-0817' | IDH wild type |
| 40 | 'TCGA-14-0865' | IDH wild type |
| 41 | 'TCGA-14-1043' | IDH wild type |
| 42 | 'TCGA-14-1395' | IDH wild type |
| 43 | 'TCGA-14-1396' | IDH wild type |

| | | |
|---|---|---|
| 44 | 'TCGA-14-1453' | IDH wild type |
| 45 | 'TCGA-19-5950' | IDH wild type |
| 46 | 'TCGA-14-1794' | IDH wild type |
| 47 | 'TCGA-14-1795' | IDH wild type |
| 48 | 'TCGA-14-1823' | IDH wild type |
| 49 | 'TCGA-14-1825' | IDH wild type |
| 50 | 'TCGA-14-1829' | IDH wild type |
| 51 | 'TCGA-02-0069' | IDH wild type |
| 52 | 'TCGA-02-0070' | IDH wild type |
| 53 | 'TCGA-02-0075' | IDH wild type |
| 54 | 'TCGA-02-0085' | IDH wild type |
| 55 | 'TCGA-14-3477' | IDH wild type |
| 56 | 'TCGA-19-2624' | IDH wild type |
| 57 | 'TCGA-19-2631' | IDH wild type |
| 58 | 'TCGA-19-5951' | IDH wild type |
| 59 | 'TCGA-19-5953' | IDH wild type |
| 60 | 'TCGA-CS-6668' | IDH mutated |
| 61 | 'TCGA-CS-6669' | IDH wild type |
| 62 | 'TCGA-19-5958' | IDH wild type |
| 63 | 'TCGA-19-5960' | IDH wild type |
| 64 | 'TCGA-27-1830' | IDH wild type |
| 65 | 'TCGA-HT-A61B' | IDH mutated |
| 66 | 'TCGA-HT-A61A' | IDH mutated |
| 67 | 'TCGA-76-6664' | IDH wild type |
| 68 | 'TCGA-HT-8558' | IDH wild type |
| 69 | 'TCGA-HT-8564' | IDH wild type |
| 70 | 'TCGA-HT-A4DS' | IDH wild type |
| 71 | 'TCGA-06-2570' | IDH mutated |
| 72 | 'TCGA-19-5954' | IDH wild type |
| 73 | 'TCGA-HT-A5RB' | IDH mutated |
| 74 | 'TCGA-CS-5396' | IDH mutated |
| 75 | 'TCGA-HT-7860' | IDH wild type |
| 76 | 'TCGA-76-6280' | IDH wild type |
| 77 | 'TCGA-76-6662' | IDH wild type |
| 78 | 'TCGA-DU-6399' | IDH mutated |
| 79 | 'TCGA-DU-6400' | IDH mutated |
| 80 | 'TCGA-DU-6401' | IDH mutated |
| 81 | 'TCGA-DU-6404' | IDH wild type |
| 82 | 'TCGA-DU-8158' | IDH wild type |
| 83 | 'TCGA-02-0003' | IDH wild type |
| 84 | 'TCGA-02-0006' | IDH wild type |
| 85 | 'TCGA-02-0009' | IDH wild type |
| 86 | 'TCGA-02-0027' | IDH wild type |
| 87 | 'TCGA-DU-6408' | IDH mutated |

| | | |
|---|---|---|
| 88 | 'TCGA-DU-8162' | IDH wild type |
| 89 | 'TCGA-FG-A4MU' | IDH wild type |
| 90 | 'TCGA-DU-A5TY' | IDH wild type |
| 91 | 'TCGA-06-0119' | IDH wild type |
| 92 | 'TCGA-DU-6407' | IDH mutated |
| 93 | 'TCGA-DU-6410' | IDH mutated |
| 94 | 'TCGA-02-0064' | IDH wild type |
| 95 | 'TCGA-HT-7879' | IDH mutated |
| 96 | 'TCGA-HT-7606' | IDH mutated |
| 97 | 'TCGA-HT-7608' | IDH mutated |
| 98 | 'TCGA-FG-A87N' | IDH mutated |
| 99 | 'TCGA-HT-7468' | IDH mutated |
| 100 | 'TCGA-HT-7471' | IDH mutated |
| 101 | 'TCGA-HT-7476' | IDH mutated |
| 102 | 'TCGA-HT-7478' | IDH mutated |
| 103 | 'TCGA-HT-7481' | IDH mutated |
| 104 | 'TCGA-HT-7603' | IDH mutated |
| 105 | 'TCGA-HT-7686' | IDH mutated |
| 106 | 'TCGA-HT-7690' | IDH mutated |
| 107 | 'TCGA-76-6285' | IDH wild type |
| 108 | 'TCGA-76-6286' | IDH wild type |
| 109 | 'TCGA-02-0060' | IDH wild type |
| 110 | 'TCGA-02-0068' | IDH wild type |
| 111 | 'TCGA-HT-7610' | IDH mutated |
| 112 | 'TCGA-HT-7677' | IDH mutated |
| 113 | 'TCGA-HT-7880' | IDH mutated |
| 114 | 'TCGA-HT-7902' | IDH mutated |
| 115 | 'TCGA-HT-8010' | IDH mutated |
| 116 | 'TCGA-HT-7695' | IDH mutated |
| 117 | 'TCGA-HT-8108' | IDH mutated |
| 118 | 'TCGA-HT-8110' | IDH wild type |
| 119 | 'TCGA-HT-7882' | IDH wild type |
| 120 | 'TCGA-76-6656' | IDH wild type |
| 121 | 'TCGA-02-0011' | IDH wild type |
| 122 | 'TCGA-76-6657' | IDH wild type |
| 123 | 'TCGA-DU-6405' | IDH wild type |
| 124 | 'TCGA-HT-8012' | IDH mutated |
| 125 | 'TCGA-HT-A5R7' | IDH mutated |
| 126 | 'TCGA-HT-8013' | IDH mutated |
| 127 | 'TCGA-HT-A615' | IDH mutated |
| 128 | 'TCGA-06-0238' | IDH wild type |
| 129 | 'TCGA-06-0240' | IDH wild type |
| 130 | 'TCGA-DU-5852' | IDH wild type |
| 131 | 'TCGA-HT-A616' | IDH mutated |

| | | |
|---|---|---|
| 132 | 'TCGA-DU-6542' | IDH mutated |
| 133 | 'TCGA-FG-7643' | IDH wild type |
| 134 | 'TCGA-HT-7605' | IDH mutated |
| 135 | 'TCGA-HT-7681' | IDH mutated |
| 136 | 'TCGA-FG-A4MT' | IDH mutated |
| 137 | 'TCGA-DU-7298' | IDH mutated |
| 138 | 'TCGA-DU-A5TT' | IDH wild type |
| 139 | 'TCGA-DU-7015' | IDH mutated |
| 140 | 'TCGA-DU-8165' | IDH wild type |
| 141 | 'TCGA-FG-8186' | IDH mutated |
| 142 | 'TCGA-06-0122' | IDH wild type |
| 143 | 'TCGA-76-4935' | IDH wild type |
| 144 | 'TCGA-HT-A617' | IDH wild type |
| 145 | 'TCGA-FG-7637' | IDH mutated |
| 146 | 'TCGA-FG-8189' | IDH mutated |
| 147 | 'TCGA-12-1598' | IDH wild type |
| 148 | 'TCGA-12-3650' | IDH wild type |
| 149 | 'TCGA-DU-7010' | IDH mutated |
| 150 | 'TCGA-14-0789' | IDH wild type |
| 151 | 'TCGA-14-1458' | IDH mutated |
| 152 | 'TCGA-14-1821' | IDH mutated |
| 153 | 'TCGA-DU-7008' | IDH mutated |
| 154 | 'TCGA-14-0813' | IDH wild type |
| 155 | 'TCGA-DU-5855' | IDH mutated |
| 156 | 'TCGA-14-1456' | IDH mutated |
| 157 | 'TCGA-CS-5390' | IDH mutated |
| 158 | 'TCGA-CS-5393' | IDH mutated |
| 159 | 'TCGA-CS-5394' | IDH mutated |
| 160 | 'TCGA-CS-5395' | IDH wild type |
| 161 | 'TCGA-27-1835' | IDH wild type |
| 162 | 'TCGA-12-0773' | IDH wild type |
| 163 | 'TCGA-12-0775' | IDH wild type |
| 164 | 'TCGA-12-0829' | IDH wild type |
| 165 | 'TCGA-76-4925' | IDH wild type |
| 166 | 'TCGA-CS-4938' | IDH mutated |
| 167 | 'TCGA-CS-4941' | IDH wild type |
| 168 | 'TCGA-CS-4942' | IDH mutated |
| 169 | 'TCGA-CS-4943' | IDH mutated |
| 170 | 'TCGA-CS-4944' | IDH mutated |
| 171 | 'TCGA-DU-7013' | IDH wild type |
| 172 | 'TCGA-DU-7018' | IDH mutated |
| 173 | 'TCGA-DU-7019' | IDH mutated |
| 174 | 'TCGA-DU-7294' | IDH mutated |
| 175 | 'TCGA-DU-7299' | IDH mutated |

| | | |
|---|---|---|
| 176 | 'TCGA-DU-7300' | IDH mutated |
| 177 | 'TCGA-DU-7302' | IDH mutated |
| 178 | 'TCGA-DU-7304' | IDH mutated |
| 179 | 'TCGA-DU-7306' | IDH mutated |
| 180 | 'TCGA-DU-7309' | IDH mutated |
| 181 | 'TCGA-DU-8163' | IDH mutated |
| 182 | 'TCGA-DU-8164' | IDH mutated |
| 183 | 'TCGA-DU-8166' | IDH mutated |
| 184 | 'TCGA-DU-8167' | IDH mutated |
| 185 | 'TCGA-DU-8168' | IDH mutated |
| 186 | 'TCGA-DU-A5TP' | IDH mutated |
| 187 | 'TCGA-DU-A5TR' | IDH mutated |
| 188 | 'TCGA-DU-A5TW' | IDH mutated |
| 189 | 'TCGA-EZ-7264' | IDH mutated |
| 190 | 'TCGA-FG-6692' | IDH wild type |
| 191 | 'TCGA-FG-7634' | IDH mutated |
| 192 | 'TCGA-76-4926' | IDH wild type |
| 193 | 'TCGA-76-4928' | IDH wild type |
| 194 | 'TCGA-76-4931' | IDH wild type |
| 195 | 'TCGA-14-0871' | IDH wild type |
| 196 | 'TCGA-14-1034' | IDH wild type |
| 197 | 'TCGA-14-1037' | IDH wild type |
| 198 | 'TCGA-19-1388' | IDH wild type |
| 199 | 'TCGA-19-1390' | IDH wild type |
| 200 | 'TCGA-HT-8107' | IDH wild type |
| 201 | 'TCGA-76-6191' | IDH wild type |
| 202 | 'TCGA-76-6193' | IDH wild type |
| 203 | 'TCGA-76-6663' | IDH wild type |
| 204 | 'TCGA-HT-7692' | IDH mutated |
| 205 | 'TCGA-HT-7693' | IDH mutated |
| 206 | 'TCGA-HT-7694' | IDH mutated |
| 207 | 'TCGA-HT-7855' | IDH mutated |
| 208 | 'TCGA-19-1789' | IDH wild type |
| 209 | 'TCGA-19-2620' | IDH wild type |
| 210 | 'TCGA-08-0392' | IDH wild type |
| 211 | 'TCGA-CS-6186' | IDH wild type |
| 212 | 'TCGA-CS-6188' | IDH wild type |
| 213 | 'TCGA-HT-7856' | IDH mutated |
| 214 | 'TCGA-HT-A5RC' | IDH wild type |
| 215 | 'TCGA-FG-A6J1' | IDH mutated |
| 216 | 'TCGA-HT-7604' | IDH mutated |
| 217 | 'TCGA-CS-6290' | IDH mutated |
| 218 | 'TCGA-CS-6665' | IDH mutated |
| 219 | 'TCGA-CS-5397' | IDH wild type |

| | | |
|---|---|---|
| 220 | 'TCGA-HT-7875' | IDH mutated |
| 221 | 'TCGA-HT-8105' | IDH mutated |
| 222 | 'TCGA-HT-7684' | IDH mutated |
| 223 | 'TCGA-HT-8106' | IDH mutated |
| 224 | 'TCGA-HT-7877' | IDH mutated |
| 225 | 'TCGA-HT-8018' | IDH mutated |
| 226 | 'TCGA-HT-8111' | IDH mutated |
| 227 | 'TCGA-HT-8113' | IDH mutated |
| 228 | 'TCGA-HT-8114' | IDH mutated |
| 229 | 'TCGA-HT-A614' | IDH mutated |
| 230 | 'TCGA-02-0054' | IDH wild type |
| 231 | 'TCGA-HT-7472' | IDH mutated |
| 232 | 'TCGA-HT-7473' | IDH mutated |
| 233 | 'TCGA-HT-7475' | IDH mutated |
| 234 | 'TCGA-06-0139' | IDH wild type |
| 235 | 'TCGA-06-0145' | IDH wild type |
| 236 | 'TCGA-HT-A619' | IDH mutated |
| 237 | 'TCGA-06-0154' | IDH wild type |
| 238 | 'TCGA-06-0158' | IDH wild type |
| 239 | 'TCGA-06-0138' | IDH wild type |
| 240 | 'TCGA-06-0174' | IDH wild type |
| 241 | 'TCGA-06-0176' | IDH wild type |
| 242 | 'TCGA-FG-5964' | IDH mutated |
| 243 | 'TCGA-06-0189' | IDH wild type |
| 244 | 'TCGA-06-0190' | IDH wild type |
| 245 | 'TCGA-06-0192' | IDH wild type |
| 246 | 'TCGA-06-0644' | IDH wild type |
| 247 | 'TCGA-06-0646' | IDH wild type |
| 248 | 'TCGA-06-1806' | IDH wild type |
| 249 | 'TCGA-06-5413' | IDH wild type |
| 250 | 'TCGA-FG-6688' | IDH wild type |
| 251 | 'TCGA-DU-A5TU' | IDH mutated |
| 252 | 'TCGA-CS-6670' | IDH mutated |
| 253 | 'TCGA-DU-5849' | IDH mutated |
| 254 | 'TCGA-DU-5851' | IDH mutated |
| 255 | 'TCGA-DU-5853' | IDH mutated |
| 256 | 'TCGA-DU-5854' | IDH wild type |
| 257 | 'TCGA-DU-5871' | IDH mutated |
| 258 | 'TCGA-DU-5872' | IDH mutated |
| 259 | 'TCGA-DU-5874' | IDH mutated |
| 260 | 'TCGA-DU-6395' | IDH mutated |
| 261 | 'TCGA-DU-6397' | IDH mutated |
| 262 | 'TCGA-DU-7301' | IDH mutated |
| 263 | 'TCGA-FG-5963' | IDH wild type |

| | | |
|---|---|---|
| 264 | 'TCGA-HT-7854' | IDH wild type |
| 265 | 'TCGA-HT-8019' | IDH wild type |
| 266 | 'TCGA-02-0116' | IDH wild type |
